# Once upon a time in Mexico: Holocene phylogeography of the spotted bat (*Euderma maculatum*)

**DOI:** 10.1101/2022.08.26.505484

**Authors:** Daniel E. Sanchez, Faith M. Walker, Colin J. Sobek, Cori Lausen, Carol L. Chambers

## Abstract

Holocene-era range expansions are relevant to understanding how a species might respond to the warming and drying climates of today. The harsh conditions of North American deserts have phylogenetically structured desert bat communities but differences in flight capabilities are expected to affect their ability to compete, locate, and use habitat in the face of modern climate change. A highly vagile but data-deficient bat species, the spotted bat (*Euderma maculatum*) is thought to have expanded its range from central Mexico to western Canada during the Holocene. With specimens spanning this latitudinal extent, we coupled phylogeography (mtDNA) with ecological niche modeling (ENM) to investigate the Holocene biogeography from the rear to leading edges. The ENM and phylogeny supported a Holocene range expansion from Mexico with increased expansion throughout the intermountain west within the last 6 kya. Long-term isolation at the southern-most margin of the range suggests one or more populations were left behind as climate space contracted and are currently of unknown status. The species appears historically suited to track shifts in climate space but differences in flight behaviors between leading edge and core-range lineages suggest that range expansions could be influenced by differences in habitat quality or climate (e.g., drought). Although its vagility could facilitate the tracking of environmental change and thereby extinction avoidance, anthropogenic pressures at the core range could still threaten the ability for beneficial alleles to expand into the leading edge.

## Introduction

Modern-day climate change is an existing and threatening influence on the distribution of biodiversity (1,2). Western North America is becoming warmer and drier (3,4), a trend that is already shifting the trajectories of ecosystems in North America (5). Globally, species are responding to warming climates by expanding into northern latitudes or into higher elevations (6). Species unable to expand their ranges must rely on phenotypic plasticity or retain enough genetic diversity to adapt to changes in their environment (7,8). The more prominent effects of climate change occur on the leading and rear edges of a species distribution (9), which are often observed as a latitudinal (polar) orientation in North America. Leading edge populations are positioned on the frontline of environmental change and may live in the colder extremes of their physiology (10). Rear edge populations may be either stable or trailing (11). Trailing rear edge populations endure warmer extremes and are likely to face extirpation. Stable rear edge populations persist in the warmest extremes and are substantially isolated from the core distribution. Increasingly, models are built to predict future distributions of a species by using contemporary geographic occurrences (12). Yet suitability maps of shifting climate alone do not provide insight into how the species might respond to those shifts. The events of the Pleistocene and Holocene provide the most recent frame of reference for extreme climate change and phylogeographic analysis at the transition of those eras can provide substantial insight into how a species may respond in the future (13).

Bats are remarkable indicators of climate change due to their powered flight (14). In recent decades, a common and relatively sedentary bat of the Mediterranean, Kuhl’s pipistrelle (*Pipistrellus kuhlii*), has consistently expanded its range into northesastern Europe in response to warming winter temperatures in the upper latitudes (15). Bats of the harsh deserts of western North America provide a relevant arena of study for range expansions. Over interspecific time scales these bat communities are thought to have been structured by habitat filtering of their harsh, arid environments (16). However, their ability to compete, locate, and use habitat warming and drying climate of today may vary by their species-specific dispersal characteristics (17). In general, species capable of shifting their range with changing climates can avoid extinction (18) and long distance dispersal is expected to play a critical role toward that end (19).

The spotted bat (*Euderma maculatum*) exhibits long flight capabilities and large home range sizes relative to many Vesper bats in North America (20), and has a poorly understood biogeographic legacy. Its three white, dorsal spots, large ears, and an audible, low-frequency call are charismatic hallmarks for identification (21–23). However, this charismatic bat is rarely observed. The species occupies a patchy, widespread range, and is associated with rugged environments from central Mexico to British Columbia (BC), Canada (22,24,25) (Fig 1). The species largely occupies desert and xeric shrubland biome but individuals at the leading edge occupy an area broadly associated with the temperate coniferous forest biome (26) (Fig 1). The space used by individuals on the leading edge, however, is semi-arid, resembling elements of habitat in the core range (23). In British Columbia, the northern-most extent of its range, *E. maculatum* is associated with hot dry river valleys comprised of bunchgrass, sagebrush and open canopy patches of ponderosa pine (*Pinus ponderosa*) and Douglas-fir (*Pseudotsuga menziesii*) trees (27,28). During the day, *E. maculatum* roosts in sheer cliffs (20,21). At night, individuals may travel over a range of elevations and vegetation types to forage, ranging from xeric lowland, subalpine meadows and pine (*Pinus*) forests to alpine meadows and spruce-fir (*Picea-Abies*) forests (−53 m to 3230 m elev.) (22,29). In southwestern North America, *E. maculatum* is capable of engaging in long, nightly flights (≤ 70 km roundtrip) and uses ponds situated in open meadows (20,29,30). The cryptic nature of this species has led to a limited number of specimens for study (31) and information is largely from hotspot localities or where bats were found dead or dying (32). This dearth of information has led most wildlife agencies in the western United States to recognize the species as one in need of special management (33,34). Canada lists the species as one of special concern (35).

**Fig 1.**
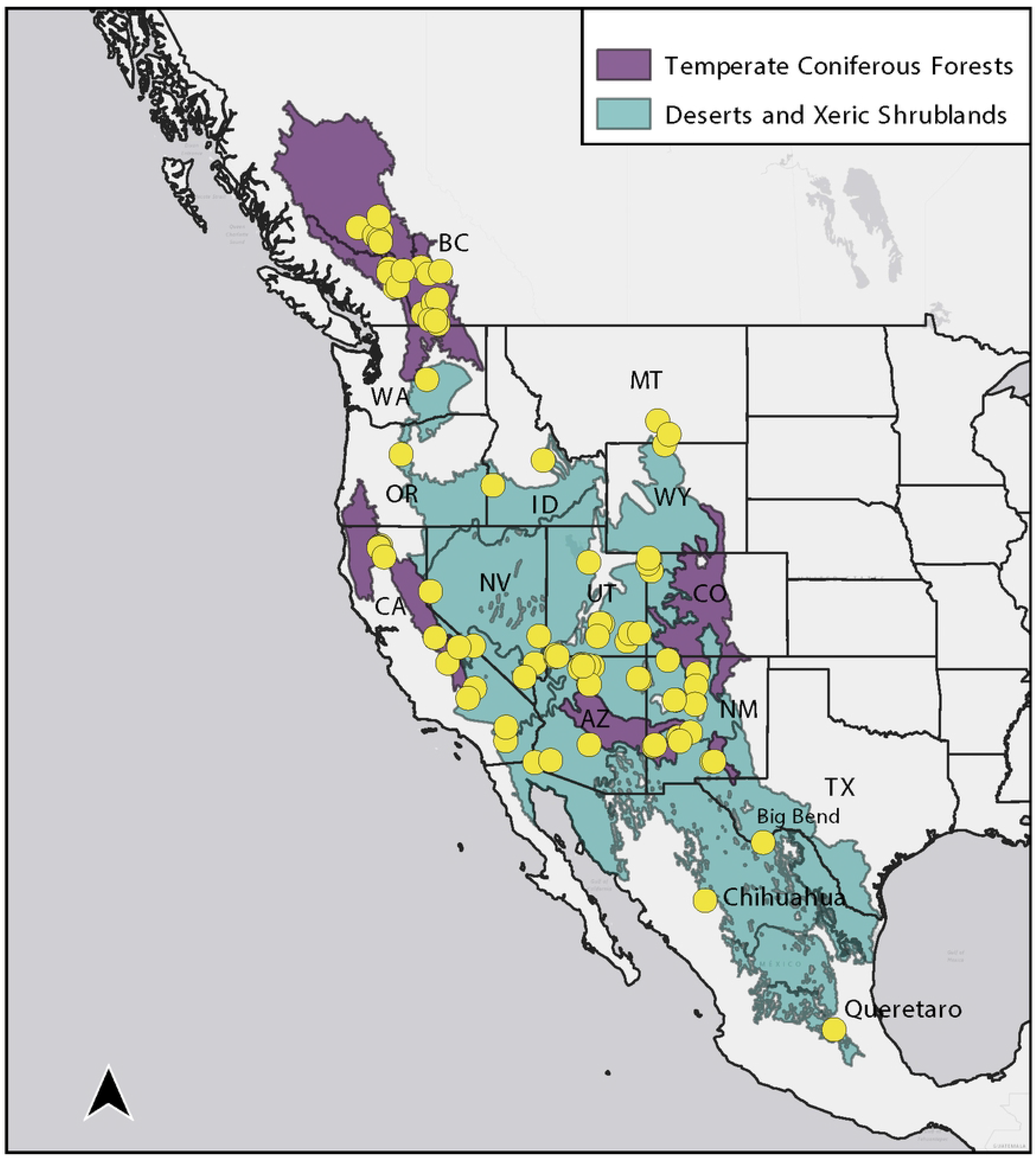
Map of historic and contemporary capture and acoustic occurrences of the spotted bat. Biomes 5 (Temperate Coniferous Forests) and 13 (Deserts and Xeric Shrublands) are derived from the World Wildlife Federation terrestrial ecosystem layers (Olson et al. 2001). States, provinces, or key localities where the spotted bat (*Euderma maculatum*) occurs are annotated. Note, that although British Columbian range is broadly categorized as temperate coniferous forest in this map, spotted bats in the region are most typically captured at lower elevation, grassland/sagebrush ecosites.

The species likely originated in Mexico and much of the climate of its contemporary range was likely inhospitable prior to the Holocene (36). This is suggested by a fossil record of a mummified specimen from the southwestern United States approximately 10.5 kya and the extant use of that locality by *E. maculatum*. An analysis of morphological traits suggested strong geographic variation (37). When geographic groups were assumed a priori, the morphologies revealed that leading edge specimens of *E. maculatum* consistently formed a distinct cluster and those of the southern-most extent, including Queretaro, formed a rooting point. Despite the geographically informative patterns, intra-geographic structure and the phylogeographic relationships of those in the core range remain unresolved.

The complementarity of phylogeography and ecological niche modeling (ENM) can further resolve the biogeographic legacy of *E. maculatum*. An ENM can assist with the interpretation of phylogenetic pattern. Phylogenetic pattern can be used to interpret the prediction of an ENM. In combination with phylogeography, hindcasted ENMs have revealed glacial refugia in *Rhinolophus ferrumequinum* in east Asia (38), and allowed interpretation of postglacial colonization of *Barbasetlla barbastellus* (39) and European *Plecotus* (40). A global analysis of bat species revealed demographic expansion of species in temperate regions and demographic bottlenecks of species in neotropical regions (41). Moreover, paleomodeling provides a way to interpret future effects of climate change with respect to addressing whether climatic tolerances have remained similar over time (40,42,43). That is, if the predicted paleo-range of a species is consistent with phylogeographically-inferred refugia, then the climatic tolerance of a species could be assumed for future scenarios (i.e., niche conservatism).

Our work seeks to address how a changing post-glacial climate might have influenced phylogeographic structure of *E. maculatum*. Spanning the entire latitudinal range of the species, we used specimen records and their mtDNA of *E. maculatum* to prime future investigations of the phylogeography and give insight into its Holocene range expansion. First, we asked whether the inferences from the *E. maculatum* fossil record provide a valid frame of reference for divergence dating (i.e., was Mexico the

Pleistocene range, with its contemporary US range unsuitable?). We examined this by modeling climate suitability from the last glacial maximum (LGM), the mid-Holocene, and contemporary climate space. Second, we asked whether maternal genetic data could better resolve phylogeographic structure in *E. maculatum* and allow us to calibrate the divergence of their lineages. To do so, we studied the mtDNA of *E. maculatum*, albeit without mtDNA from specimens representing the Mexican state of Chihuahua and the US states of California, Texas, and Idaho. To account for this data deficiency, we contrasted the mid-Holocene paleoclimate to contemporary climate space to elucidate routes of expansion. Given the latitudinal extent of the specimens, we hypothesized that mtDNA will resolve finer phylogeographic structure for leading edge (BC, CAN) and rear edge populations (Queretaro, MEX), and a core range. We examined this by resolving phylogeographic structure, phylogenetic substitutions, and by temporally calibrating the phylogeny with information from the fossil record. Third, we discuss how such a highly vagile bat species might track changes in one of the world’s regions most impacted by climate change.

## Materials and methods

### Ecological niche modeling

The context of our ENM was based on occurrences largely encompassing the core range (S1 Fig 1) and from likely foraging sites (i.e., mist-net captures). We performed all data pre-processing and spatial analyses in R v.3.2.2 (44). We assembled geographic coordinates from capture and museum specimen locations. We filtered out early records that were of *E. maculatum* found dead or dying in areas that may not represent elements of habitat for foraging and roosting (e.g., fringes of elevation range, driveways, and warehouses). We also filtered out occurrences made vague out of concern for the animals and sites. Acoustic occurrences were omitted due to differences and uncertainties between where an animal was traveling and foraging. Under our target context, a single occurrence remained at one of the most northern portions of the range (Lillooet, BC, CAN). However, this occurrence was omitted because it could be due to finer-scale temporal variability at the leading edge and therefore may not represent its historic climate tolerance (45). Under this setting, occurrences may still exhibit biases because they may represent hotspots for *E. maculatum* activity or preferred capture locations so we dereplicated occurrences within 1km^2^ grid cells. Finally, we retained occurrences after 1970 to match the range of years used to generate bioclimatic variables (46), resulting in 40 distinct occurrences for modelling the ENM.

To build the ENM, we first selected all bioclim variables as potential predictors at 30 arc second resolution (46). To avoid multi-collinearity among predictors in training a model, we extracted and Z-transformed raster values of the occurrences and conducted an analysis of principal components using the Pysch package (47). We then selected representative predictors from 4 principal components: mean annual temperature (bio1), mean diurnal range (bio2), isothermality (bio3), and precipitation coldest quarter (bio19). The predictors were selected toward the goal of predicting spatial pattern as opposed to interpretation of environmental parameters. We modeled a presence-only ecological niche using Maxent v3.3.3 (48) in the biomod2 package (49). Maxent models are trained by minimizing the relative entropy between probability densities of species occurrence and environmental background in covariate space (50). Maxent is the most widely used algorithm for prediction because it was developed to model presence-only records. The theoretical limitations and cautions of Maxent have been well documented (51,52) but Maxent can generate useful models for rare species, particularly those with 25-50 occurrences or more (53). The biomod2 package provided a platform for conducting the Maxent modeling as well as combining independent model runs for consensus projection (i.e., ensembling). To pull background points, we generated a training background distribution calculated from a 1σ buffer from means of each occurrence value (54) using altitude (alt) (46), bio2, bio3, and bio19. This allowed us to generate a background that exceeded the constraints of the occurrences to provide more flexibility in generating background points (S1 Fig 1). We trained Maxent models with an 80:20 split of the occurrences for training and testing. We used three sets of randomly sampled background points (n = 120 per set) and ran the model through 10 evaluations. Each set of evaluations for a model were then combined into a full model. We evaluated the models using the area under the receiving operating curve score for training and test sets (AUC_train_, AUC_test_) and the true skill statistic (TSS). For ensembling, we included any full model with AUC_test_ > 0.7 for projection into contemporary, mid-Holocene (~ 6 Kya), and last glacial maximum (~ 22 Kya) climate space. We used the TSS as a binary threshold to provide balance between sensitivity and specificity for contrasting gained, retained, and receded climate space.

### Genetic sampling and mtDNA sequencing

We acquired 27 *E. maculatum* tissues and 1 *Idionycteris phyllotis* tissue from live capture and museum specimens that covered the entire known latitudinal range of the focal species (Table 1). We captured individuals between 2009 and 2014 using mist nets and collected wing biopsies and buccal swabs with the approval of the Northern Arizona University Institutional Animal Care and Use Committee (07-006, 07-006-R1, 07-006-R2; Walker et al., 2014); in Canada, wing biopsies were collected under a British Columbia research permit (KA14-148262-1). We handled bats following the guidelines of the American Society of Mammalogists (55). Museum tissues were skin clips excised from the midline of specimens originally collected between 1981 and 2013. We extracted genomic DNA from wing biopsies of live-captured individuals using the DNeasy Blood and Tissue protocol (Qiagen, Valencia, CA, USA). DNA from museum samples was extracted using previously described phenol-chloroform methods (56). We amplified a 193 bp fragment of the D-loop region using custom primers (F: Dloop_196bp, GATGCTTGGACTCAACACTG; R: RevL16517_600, GTCCTGTAACCATTAAGTTCAC). PCRs contained 2 μL gDNA, 1 μL 10X Mg-free PCR buffer (Invitrogen, Thermo Fisher Scientific, Waltham, MA, USA), 2 mM MgCl_2_, 0.2 mM of each dNTP, 0.3 U/μL Platinum Taq polymerase, and 0.5 μM of each primer in a 10 μL reaction. Thermal cycling conditions involved a 6 min denaturation at 95 °C followed by 45 cycles of denaturation at 95 °C for 30 s, annealing at 58 °C for 30 s, and polymerase extension at 72 °C for 30 s, with a final extension at 72 °C for 10 min. We also amplified a 596 bp fragment of cytochrome *b* (cytB) using custom primers (F: EUMA_Cytb_190f, GCTCCGTAGCCCACATTTGC; R: EUMA_Cytb_1027r, TGGCTGTCCAATTCAGG). These PCRs contained 2 μL gDNA, 2.5 μL 10X Mg-free PCR buffer (Invitrogen, Thermo Fisher Scientific, Waltham, MA, USA), 3 mM MgCl2, 0.2 mM of each dNTP, 0.1 U/μL Platinum Taq polymerase, 0.02 μg/μL of Bovine Serum Albumin, 0.5 μM of each primer in a 10 μL reaction. Cycling conditions were initial denaturation of 95 °C for 10min, then 40 cycles of denaturation at 95 °C for 30 s, annealing at 62 °C for 30 s, extension at 72 °C for 2 min, and a final extension of 72°C for 10 min. Amplicon was purified using the ExoSAP-IT cleanup protocol (Affymetrix, Santa Clara, CA, USA) and subsequently Sanger-sequenced using BigDye v.3.1 Terminator kit according to the recommended protocol of the manufacturer (Applied Biosystems, Foster City, CA, USA) on an ABI3130 Genetic Analyzer (Applied Biosystems, Foster City, CA, USA). We used MJ Research PTC-200 thermal cyclers for all PCRs. Sequences were edited using Sequencher v5.3 (http://www.genecodes.com).

**Table 1.**
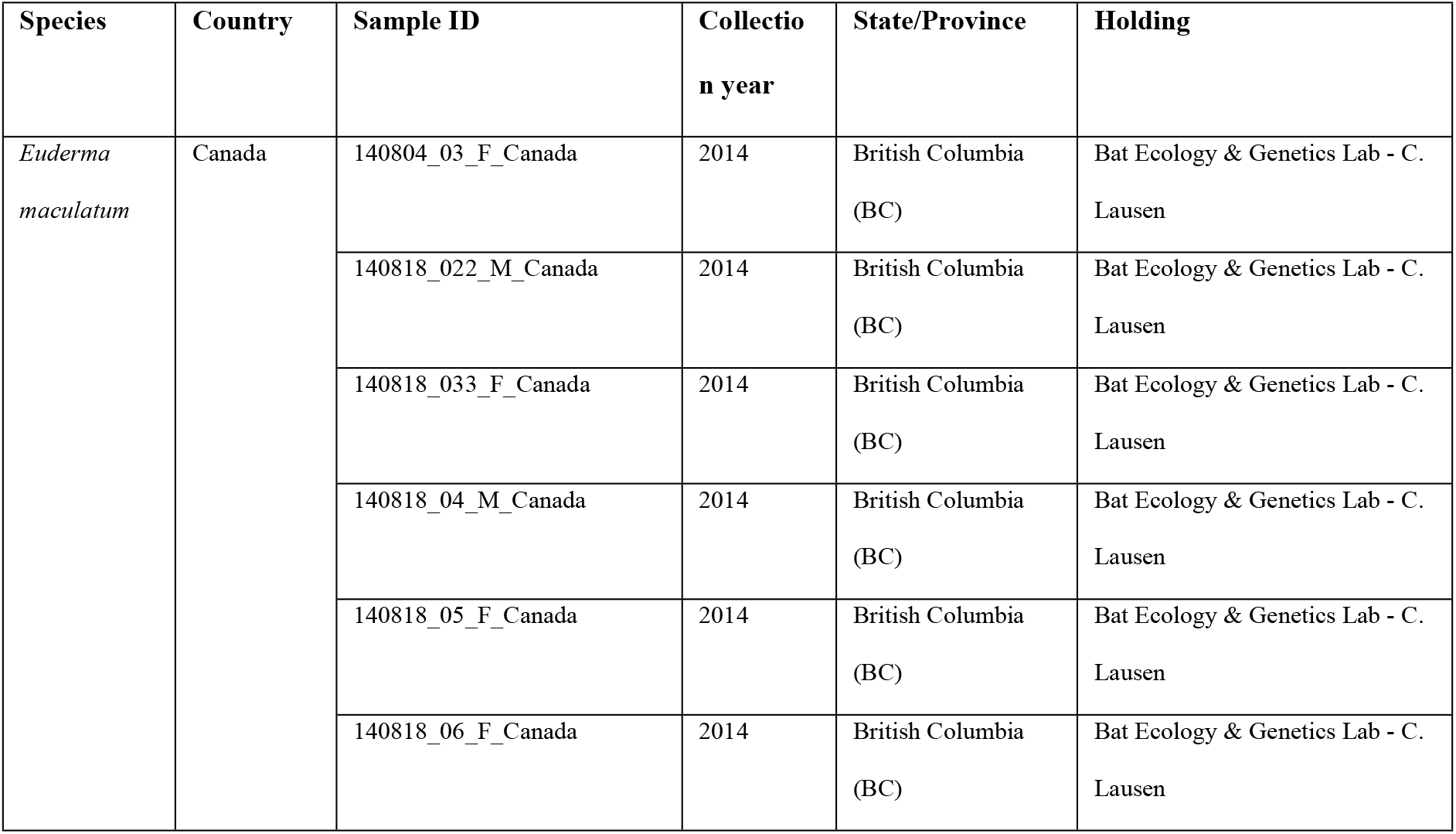
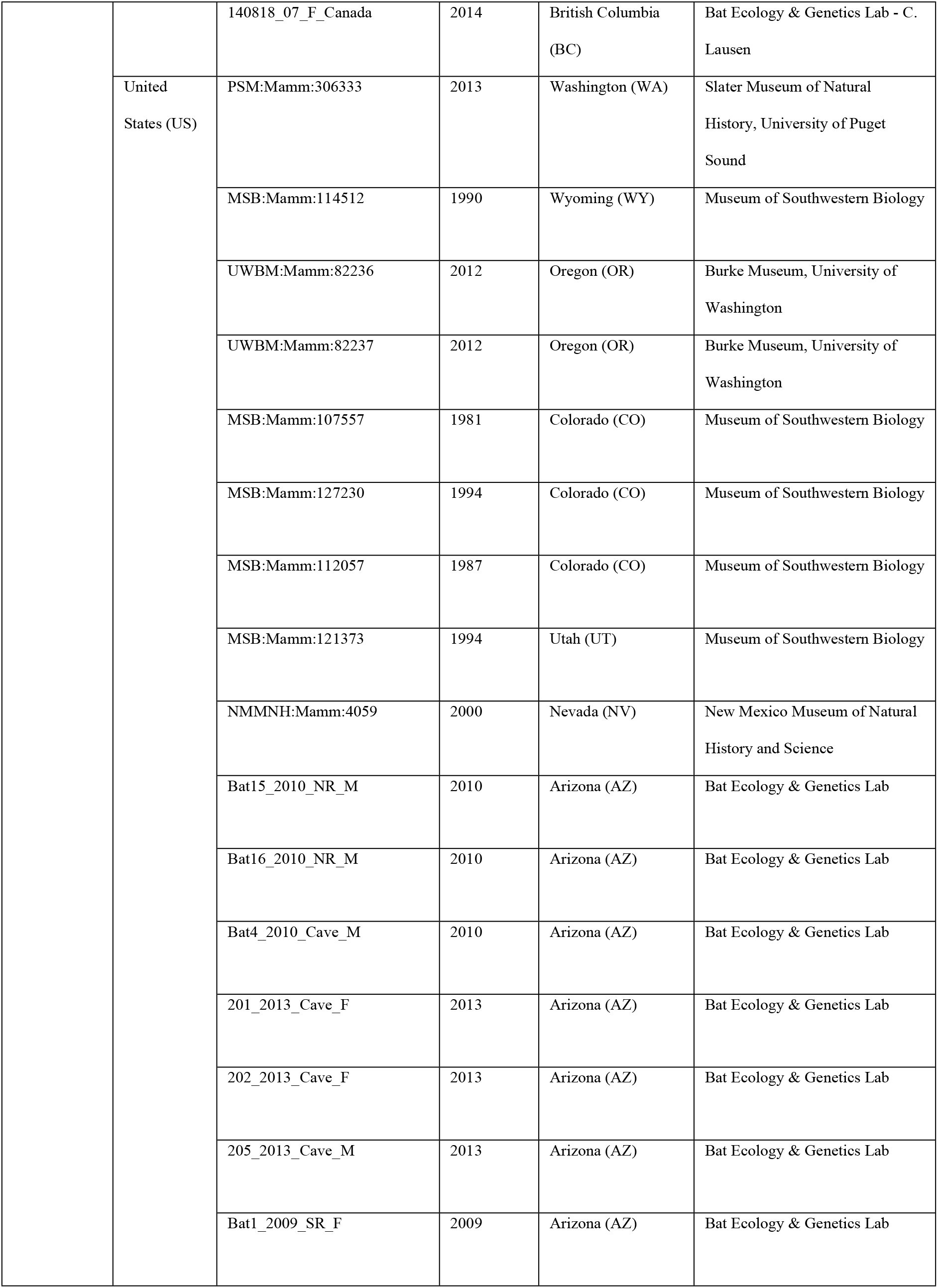
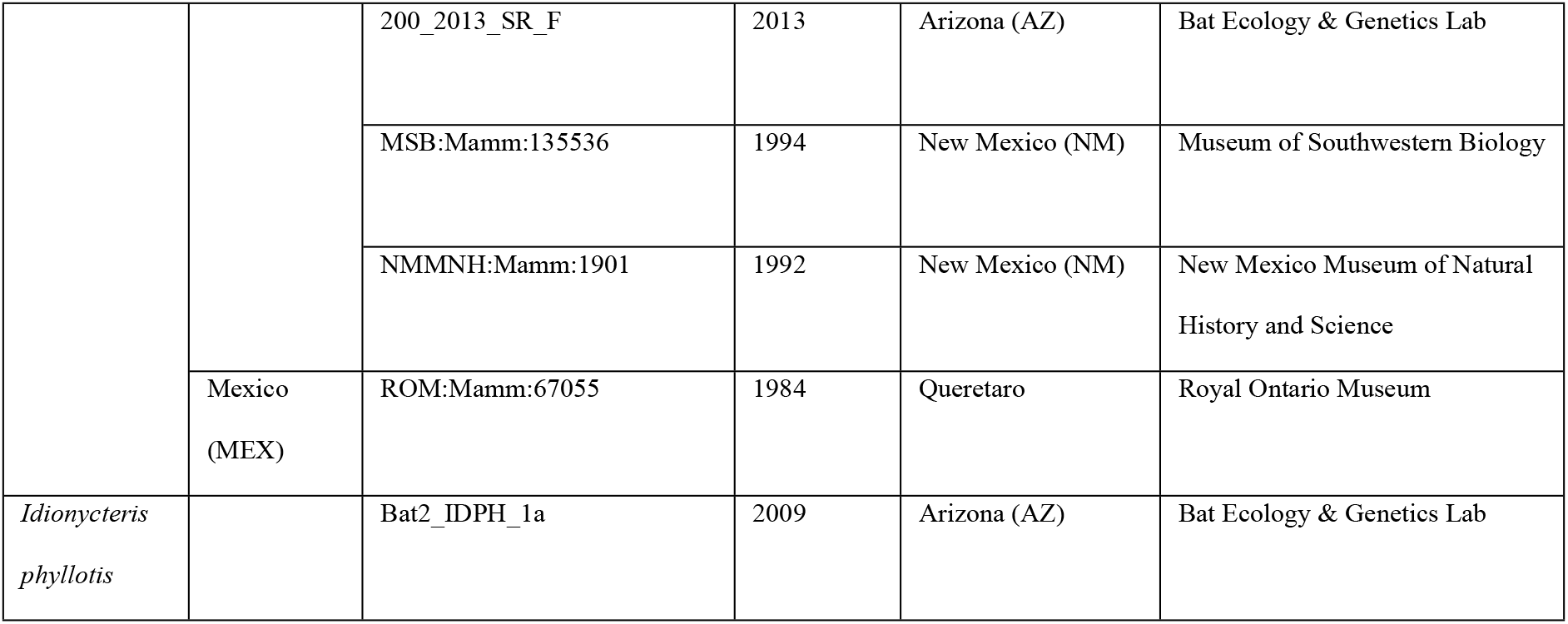
Specimens used for phylogenetic analysis and their geographic origin. Any specimen with a holding under the Bat ecology & Genetics Lab indicates an individual that was genetically sampled in our field surveys.

### Phylogenetic analysis and divergence dating

We aligned sequences of each marker using ClustalW (57) in MEGA7 (58). We concatenated both markers (total positions = 789) with substitution models estimated for each marker using BIC-based model selection in PartitionFinder v1.1.1 (59), whereby cytB was partitioned at all three codon positions. We used RAxML v8.2.8 (60) with 1000 bootstrap iterations to calculate a maximum likelihood (ML) phylogeny using the GTRGAMMA substitution model.

We estimated divergence dates with Bayesian and maximum likelihood frameworks using the topology of the ML phylogeny. We initially tested whether the markers exhibited a strict molecular clock to validate the use of relaxed molecular clocks as a prior in the Bayesian framework. Using the TRN+I substitution model, estimated as the most parsimonious from PartitionFinder, we tested whether the topology exhibited the strict molecular clock in MEGA7 using the maximum likelihood method (61), which was rejected (p < 0.0001). This result led us to proceed with divergence estimation using relaxed clock models. We first used BEAST v2.2.2 (62). We partitioned each marker using the TRN+I substitution model (TN93, Kappa1 = 2.0, Kappa2 = 2.0, Proportion Invariant = 0.5). We used a relaxed, lognormal clock model with default parameters and a coalescence tree prior that assumed constant population size (distribution = 1/X). Mean clock rate (ucldMean) was provided an exponential distribution with mean of 1.0 with standard deviation less than 1 (95% probability) under a gamma distribution. Guided by the topology of the ML tree, we constrained two clades that diverged from Mexico without parametric assumption. We calibrated the basal node of the constrained clades using the time that the oldest known specimen lived (~10.5 kya) under two parametric assumptions (calibration priors). We provided a lognormal distribution for the first model and given an offset of 10,500 years (hard minimum based on mummified specimen), a mean of 1.0, and a standard deviation of 6.17 to give a 95% probability that the divergence occurred within the last 80,000 years (soft maximum). The lognormal distribution assumed that divergence from Mexico occurred during a time frame prior to the discovery of the mummy and is considered appropriate for modeling paleontological information (63). The second model was given a normal distribution with a mean of 10,500 years (σ = 1000). This scenario assumed divergence occurred sometime before or after the earliest known specimen. The normal distribution is more appropriate for modeling divergence following biogeographic events (63), such as post-glacial climate change. For both models, tip-dates (calendar years before present) for each individual were also included as priors for additional temporal calibration. For models built on each of the calibration priors, we conducted three randomly seeded Markov-chain Monte Carlo (MCMC) runs, including an empty run to evaluate the effects of the priors. We ran each chain for 100 million iterations, sampling every 1000 trees. We used Tracer v1.7.1 (64) to evaluate chain stationarity in each run, compare their posterior probabilities, and effective sample size (ESS > 200) estimates for each parameter. We subtracted the first 30% of the chain for each run as burn-in. We compared strength of the models built on each calibration priors using marginal likelihood estimates with default parameters in PathSampler (62). Trees from each of the three test runs were combined and consensus tree annotated in LogCombiner and TreeAnnotator (maximum clade credibility) (62). We also applied maximum likelihood-based dating using the RelTime method in MEGA7 (65). This approach allowed us to drop parametric assumptions required in BEAST2 by instead using branch lengths (i.e., nucleotide substitution) to date the phylogenetic nodes (66). We used the same concatenated sequences with *I. phyllotis* as an outgroup to generate a neighbor-joining starting tree with the same substitution models as the other analyses. We calibrated the same node as for the BEAST2 analysis at 10,500 ± 500 years before present to scale the node ages. We evaluated divergence dates in FigTree v1.4.3 (67) and phylogeographic pattern using R package phytools (68).

## Results

### Ecological niche model

ENMs generated from two of three datasets of background points gave test AUC scores (Table 1 in S1 Supporting Information) of 0.81 (AUC_train_ = 0.81) and 0.83 (AUC_train_ = 0.79), indicative of informative models (Phillips & Dudík, 2008). The similar magnitudes of AUC_test_ and AUC_train_ of these models indicated that the models were not overfit. The model that we omitted from the ensemble had an AUCtest of 0.56 and AUC_train_ of 0.83, which indicated that this model was overfit and predicted poorly. Each predictor substantially contributed to the models. The average importance of the predictors was isothermality (0.60 ± 0.11 SD), mean diurnal temperature (0.49 ± 0.35), mean annual temperature (0.31 ± 0.09), and precipitation coldest quarter (0.18 ± 0.1) (Table 2 in S1 Supporting Information). Despite some variability among model performance, continuous distribution maps (Fig 1 in S1 Supporting Information) encompassed an area largely occupied by *E. maculatum* and included known locations that were omitted in the pre-processing stage.

It is preferable to present a Maxent map as a continuous gradient of relative suitability (52) (Available in S1 Supporting Information Fig 1) but we set a binary suitability threshold to better summarize general patterns of gained, receded, and retained climate space (Fig 2). The TSS was between 0.48 and 0.54 so we set the binary threshold at 0.5 to reflect a balance between sensitivity and specificity for plotting. A comparison of projections between the LGM and the mid-Holocene (~22 to 6 kya) showed substantial expansion and contraction of predicted climate space. Climate space receded from central

**Fig 2.**
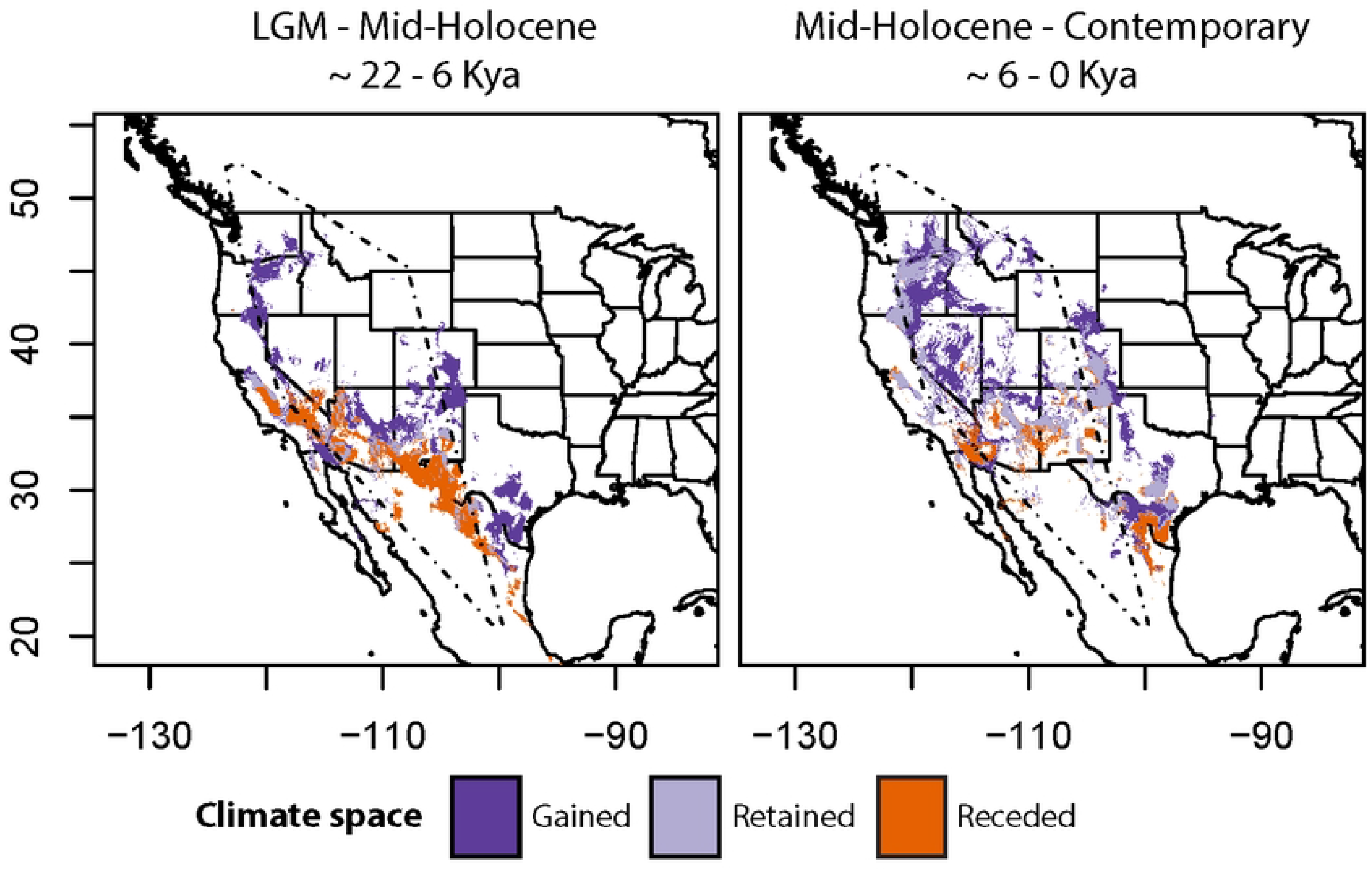
Gained, retained, and receded climate space for *E. maculatum*, predicted from a species distribution model (Maxent) (binary threshold = 0.5). Left panel: a stretch of receded climate space (in orange) spans from proximity of central Mexican specimen, approximately along the Sierra Madre Oriental to the lower southwestern United States after the transition of the last glacial maximum (LGM) to the mid-Holocene. Little of the ancestral regions were retained (in light purple) but climate space gained (in dark purple) along a narrow corridor approximately along the Sierra Nevada and Cascade mountain ranges. Right panel: Throughout the latter half of the Holocene, predicted climate space has steadily expanded as far north as British Columbia and inland into the Intermountain west. A convex polygon in dashed lines border the extent of the known species range.

Mexico into the southwestern US, whereas climate space expanded from the southwestern US in a narrow strip along the US States of CA and NV into OR and WA. A comparison of projections between the mid-Holocene and contemporary climate space (~6 to 0 kya) showed inward, patchy expansion across western North America. Patterns of receding climate space occurred in the southern range with some gained climate space approaching the Canadian range. The contemporary projection largely predicted suitable climate space within the boundaries of the known range with some under-prediction into the leading edge range of Canada and some potential over-prediction outside the eastern margins of the known *E. maculatum* range.

### Phylogenetic analysis

When rooted on the central Mexico specimen, the maximum likelihood (ML) tree (Fig 3, inset) exhibited strong phylogeographic structure. It gave a paraphyletic topology (likelihood = −1489.26), which we divided into 3 core lineages representing the rear, core, and leading edge ranges: southern, central, and northern. The southern lineage was represented by single specimen from Queretaro, Mexico. The central lineage contained specimens from southwestern and west-central US. This clade contained sub-lineages that were largely absent phylogenetic structure, except for strong clades in the US states of Colorado and Wyoming (> 80% bootstrap support). The northern lineage contained specimens from northwestern US and BC. Sub-lineages of this clade showed a deep divergence of a lineage of specimens occurring in CO and WY. More distally in the northern lineage, we also observed a shallow divergence between specimens in Oregon and the region spanning Washington and British Columbia (> 80% bootstrap support).

**Fig 3.**
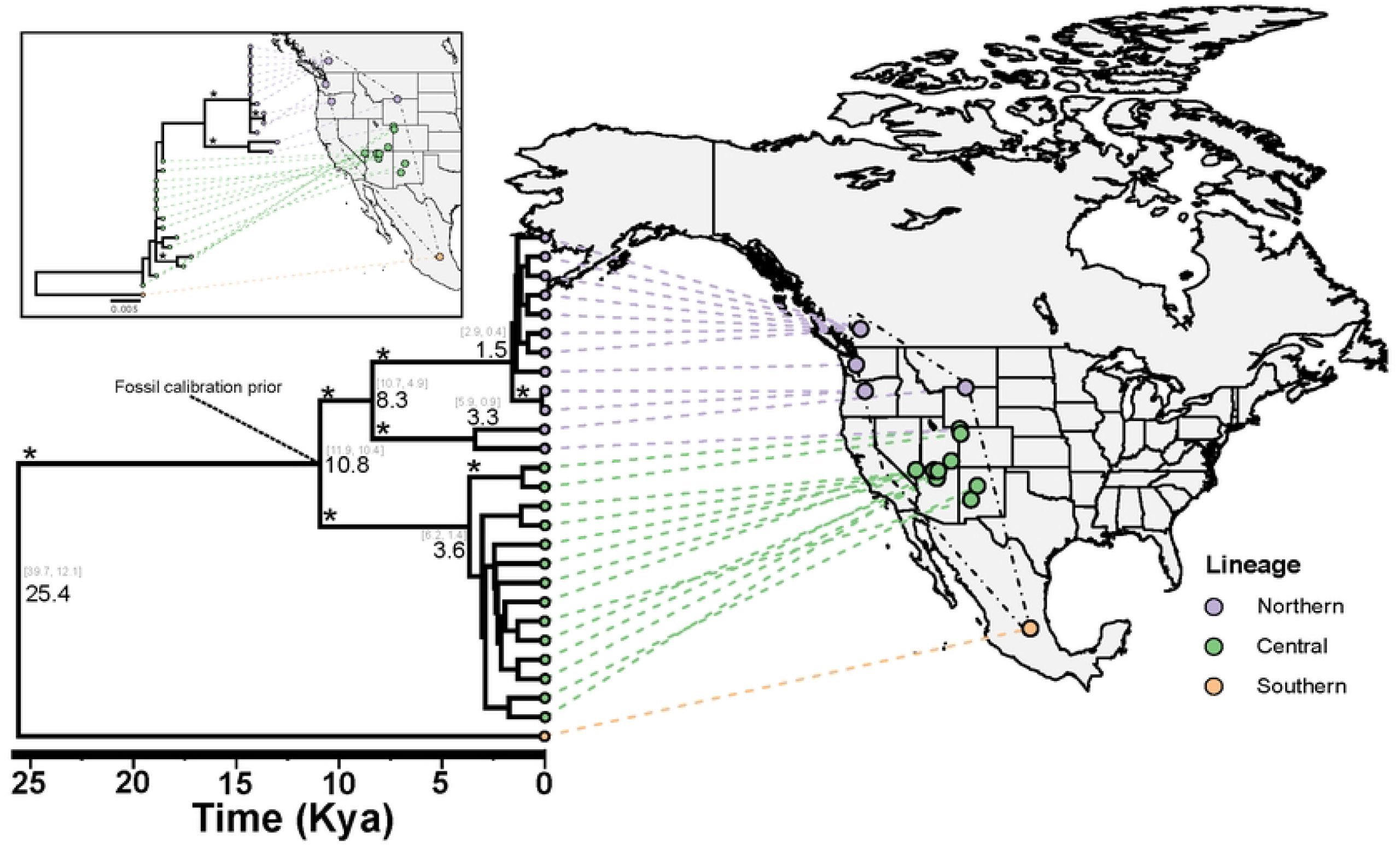
Guided by topology of ML tree (inset, scale = substitutions/site), a maximum clade credibility tree for *E. maculatum* produced from BEAST2 provides estimates of divergence times (x-axis in thousands of years, lognormal calibration shown) and 95% highest posterior density intervals (square brackets) between strong geographic lineages (* symbol indicates posterior probability >80% for BEAST2 >70% bootstrap support for ML). Assuming proper placement of calibration point, observed geographic divergence among northern-central and southern lineages began during the last glacial maximum (between 20 and 30 Kya), followed by most divergence occurring in the latter half of the Holocene (< 6 Kya). A convex polygon in dashed lines border the extent of the known species range.

We used the ML phylogeny to enforce clades for the southern, central, and northern lineages for the BEAST2 analysis (Fig 3). Under a lognormal calibration prior, two of the three MCMC chains converged with ESS > 200 but all gave identical posterior probabilities. Parameter estimates under the normal prior also converged with all runs with ESS > 200. The empty runs indicated that estimates were informed by priors and not by chance. Marginal likelihood was slightly closer to 0 for the lognormal calibration prior (−1499.56) than the normal calibration prior (−1599.33), suggesting that the models informed by the lognormal prior were stronger. Divergence estimates were largely identical among all dating priors and methods. Divergence estimates from the RelTime analysis provided higher uncertainty but divergence estimates were still contained within the 95% highest posterior density intervals estimated in BEAST2 (Fig 4). Northern and central lineages diverged from the southern lineage ~ 25 kya (Figs 1 and 2). Following assumed divergence of northern and central lineages (~ 10.8 kya), the northern lineage experienced a west-east split ~ 8 kya. Phylogenetic radiation in the southwestern US, as well as north-south splits among sub-lineages occurred within 4 kya. The most recent split occurred between populations in OR and populations in WA and BC at 1.5 kya. DNA sequences (D-loop, cytB) used for phylogenetic analysis are available in GenBank under the following accession numbers: OP234456:OP234511.

**Fig 4.**
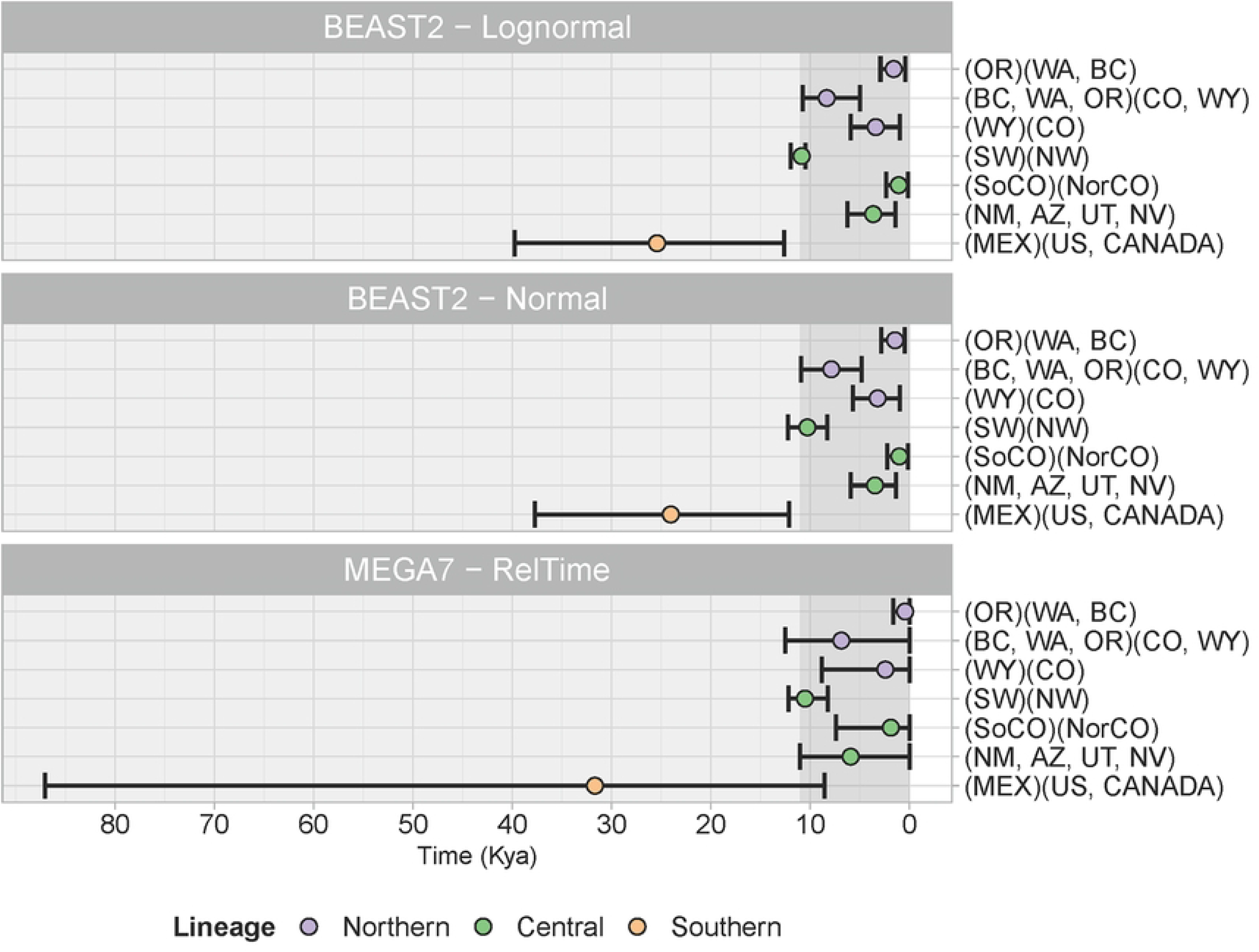
Divergence times (x-axis) among lineages were largely corroborated among calibration assumptions under Bayesian (BEAST2) and maximum likelihood frameworks (RelTime). Colored by major lineages, divergence among sub-lineages are described by abbreviations of regions (e.g, NW = northwest, SW = southwest, “No” = North, “So” = South), countries, states and provinces. Confidence intervals represent 95% highest posterior density for BEAST2 analyses and 95% confidence intervals from RelTime. Divergence is contrasted by sub-lineages within separate groups of parentheses, whereas regions within a sub-lineage are placed within a set of parentheses. The Pleistocene era is shaded in light grey and the Holocene era is shaded in dark grey.

## Discussion

In the face of unprecedented climate change, understanding the response of a species in the early Holocene is relevant to understanding its response to climate pressures in the near future. The cryptic nature of the highly vagile *E. maculatum* has led to more questions about the ecology of the species than data to address them. Despite its data deficiency, we were able to investigate its biogeographic legacy using mtDNA and ecological niche modeling. Our work supports the hypothesis of Mead & Mikesic (2001) that Mexico was the Pleistocene-era range of *E. maculatum*, later colonizing most of its US and Canadian range during the Holocene. In support of our second hypothesis, our phylogeographic evidence based on mtDNA supports, clarifies, and expands on morphologically-inferred pattern (37). The broader phylogeographic relationships reveal a stable rear edge lineage in central Mexico, a core range lineage largely throughout the US, and a leading edge lineage into BC. Our work provides a window into understanding climate-induced range extraction and contraction for a long-flying bat species in the absence of contemporary, anthropogenic pressures.

Mexico was likely the Pleistocene-era range of *E. maculatum*. This idea was originally founded on the occurrence of an early Holocene-era mummy and absence of Pleistocene-era occurrences from hotspot localities (e.g., caves and middens), ranging from the Grand Canyon to southeastern Arizona (36,69). The ENM allowed us to predict that contemporary climate space largely encompasses the mountainous periphery of the broader Great Basin (Fig 2). Nevertheless, hindcasting into the LGM indicates that climate space encompassed the mountainous regions of the Mojave and Chihuahua desert regions (ecoregions visualized in Patrick & Stephens, 2014), mostly along the Sierra Madre Occidental. We were able to predict climate space into the Queretaro locality (the southern-most margin of the range) despite training the ENM without the Queretaro occurrence, which lends strength to the generalizability of the ENM. Phylogeographic evidence shows that the southern lineage (Queretaro specimen) lies paraphyletic to all other individuals and exhibits a high amount of substitutions per site. This pattern indicates long-term isolation and supports Mexico as an ancestral range, given its geographic proximity. Assuming the fossil calibration, this matrilineal lineage approached divergence during the LGM. The consistency between this isolated lineage, prediction of niche space into that area during the LGM, and inference from the fossil record provide some support of niche conservatism in *E. maculatum* throughout the last 20,000 years. The LGM divergence at the root node and the presence of suitable climate space from central Mexico into the US may also indicate that a more northerly population in Mexico could have already been staged for expansion into the contemporary range. In particular, some of the predicted climate space during the LGM overlapped the Cuatro Ciénegas Basin, a well-known glacial refugium (70) and where *E. maculatum* has been detected (71). Coincidently, the refugium is just southeast of Big Bend, TX (Fig 1), which has been suggested as a “hotspot” locality for *E. maculatum* (22). Although specimens from Cuatro Ciénegas Basin and Big Bend localities were unavailable for phylogenetic analysis, we hypothesize that if this area was a refugium prior to expanding into the contemporary range, individuals from these localities should lie paraphyletic to the central lineage. This hypothetical lineage could provide further insight into niche conservatism.

The contemporary distribution was likely a consequence of mid-Holocene warming. The most superficial nodes and shallow radiations in the ML phylogeny (Fig 3 inset) indicate recent geographic expansions throughout the southwestern US, as well as recent colonization into the leading edge range. Phylogeographic divergence into the northeastern range (WY, CO, MO) is mirrored in the northern and central lineages. That is, this region appeared to have been colonized with subsequent divergence within both the northern and central lineages (Figs 3 and 4) at roughly the same time (~3 kya). Supported by the ENM, this coincident expansion throughout the Great Basin was likely a climate-induced phenomenon. Co-occurrence of individuals from both the northern and central lineages provides some evidence that admixture among leading and core range lineages could be occurring, albeit this phenomenon would be better addressed by nuDNA. Such zones of admixture could be important in facilitating the exchange of local, climate-adapted genes in the midst of ongoing climate change (10,72). Assuming the fossil calibration, we estimate that the superficial nodes diverged within the last 3-6 thousand years with the leading edge of the distribution in WA and BC being the youngest (1.5 kya; between 0.4 and 3 kya). This leading-edge population exhibits strong monophyly with a shallow but strong divergence from OR, indicating that the expansion into Okanagan and Fraser valleys in BC (summarized in COSEWIC 2014), Canada was relatively recent. The absence of suitable climate space in BC also provides evidence of the recent colonization. The ENM underpredicted range into BC but we believe this could be because species on the leading edge might persist in fine-scale, temporally variable, microclimate conditions (45). The lack of prediction space at the leading edge could also be explained by only having trained the ENM using central range occurrences. However, this underprediction was minimal given the close proximity of predicted suitable climate space to known range in BC. Although mtDNA and occurrences for TX, northern CA, and ID specimens were unavailable in the current study, we were able to leverage the ENM to reveal a potential route of early colonization through the Sierra Nevada and Cascade Mountain ranges (Fig 2). This is based on the early divergence between northern and central lineages. *E. maculatum* specimens are rare from CA but are known to occur along the two mountain chains from acoustic data (25). Subsequent Holocene warming likely allowed the species to expand its range inwardly throughout the Great Basin within the last six thousand years.

Much uncertainty still surrounds the timing of the divergence between the central and northern lineage because it was assumed from the fossil record and its placement was based on a partial phylogeny. Ancient DNA from the 10.5 kya mummified specimen and others that have been collected at the same locality will likely provide a more accurate form of calibration for the phylogeny. Until ancient DNA can be included in a future analysis, we recommend some caution be taken in the interpretation of the absolute times presented here. However, we believe that much of the divergence likely occurred within the temporal realm of the Holocene. The absence of climate space mostly north of Mexico provides support for the temporal assumption because all individuals in that clade occupy geographic space north of the predicted climate space of the LGM. Further support of the calibration is in the coincident expansion of suitable climate space between the LGM and the mid-Holocene.

*E. maculatum* likely owes its ability to track climate-induced change in part to its exceptional flight capabilities; however, its powered flight may be both the exception and the rule. We emphasize that our study represents an example of the historic range expansion of a western North American bat with great dispersal capabilities. Previous phylogeographic studies of North American desert bats have largely focused on species with relatively low vagility and were largely based on interspecific divergence throughout the Pleistocene (73–75). Yet it is still unclear if or how the dispersal ability of *E. maculatum* influenced the rate of poleward expansion throughout the Holocene. The phenomenon and rate of range expansion depends on numerous factors in addition to dispersal capability. Leading edge populations, like populations of *E. maculatum* in BC, are primarily expected to influence the rate of future range shifts (76). Low numbers of reproductive individuals with or without sex-bias may slow the rate of expansion and these individuals may benefit from reduced resource competition (77). There is a key contrast of dispersal between the leading edge and core range populations of *E. maculatum*. Individuals in the core range appear to use their flight for travel to foraging areas, whereas those at the leading edge maximize their flight for foraging locally. In the high elevation semi-deserts of the core population, individuals regularly returned to a known foraging site, traveling 77 km (round trip), and had an average home range of roughly 300 km^2^ (20,30). In the leading edge (Fraser and Okanagan valleys, BC), individuals also tended to return to the same local foraging sites (23,78) but were observed roosting much closer to their foraging sites (6-10 km) (78,79). Although less distance is traveled in the leading edge range, the long flight is instead leveraged for foraging in continuous flight. Hobbs et al. (78) determined that *E. maculatum* was foraging over a small area, with home ranges estimated at 1.57 km^2^ −4.58 km^2^; using radiotracking, maximum straight-line distances covered by foraging bats ranged from 6.9 to 18.8 km, with a maximum linear distance covered during a 30 minute foraging time of less than 3 km. The smaller home ranges on the leading edge could be a result of higher prey densities, lower interspecific competition, or the shorter nights in northern areas limiting foraging times (80). Dispersal-promoting behaviors of individuals on the leading edge requires further investigation. An important aspect of predicting the future response of this species, with respect to range expansion, is by further comparing dispersal characteristics, traits, and behavioral tradeoffs of leading and core range lineages. Dispersal-promoting morphologies can potentially be discovered given the high variability of quantitative traits among *E. maculatum*, even with small sample sizes (37).

Given the phylogeographic structure and differences in flight behaviors between core and leading edge ranges, we hypothesize that the species might engage in dispersal-promoting behavior in response to lower habitat quality (76) (e.g., drought, higher temperature). The greater home ranges of those in the core range could be a result of poorer habitat quality in a drier, semi-desert southwest, whereby increased travel could reflect lower selectivity in foraging site or prey selection (81,82) than individuals in the leading edge. Increasing temperature can lead to roost abandonment in female bats (83) and physiological responses to heat have been documented in the core range of *E. maculatum* at 30 °C (21). Drought in desert bats makes lactating female bats thirstier, expending more energy to drink (84) and can lower reproductive output (85). This reduction in habitat quality could increase inter and intra-specific competition at foraging sites, decrease vegetation for insect prey, decrease water availability, and therefore promote increased rates of founding new localities. During a drought in 2006, *E. maculatum* was detected at 6 new foraging localities in New Mexico (core range) and was thought to be a phenomenon of the individuals flying further distances to drink (32). Although, *E. maculatum* might be able to track new climate space, a key limiting factor is the predominant selection of fixed roost structures (e.g., sheer cliffs, crevices), which will remain despite changing climate and vegetation (86).

Although the geographic structure of *E. maculatum* can be further refined by nuclear DNA, the lineages of the rear, leading edge, and core range provide a first step for which to discuss intra-population conservation implications. At the rear edge, the southern lineage (Queretaro) may meet the criteria of a climate relict (87), stable rear edge population and warrants further survey. Populations of the southern lineage appear to be the most geographically isolated and ancient. The age of the lineage is consistent with the expectation that a stable rear edge population is two to three orders of magnitude older than lineages of the rest of the range (11). The entire range of the species may be functionally misleading due to the lack of suitable climate space into central Mexico and in the patchiness throughout Chihuahua and Big Bend regions. Additionally, we are unaware of any detections in Queretaro since 1984 (88). This population of the southern lineage could be considered an evolutionary significant unit (ESU). The definition of ESUs have varied over the years (89) but we believe that the degree of morphological clustering (37) and evidence of long-term genetic isolation, geographic isolation, differences in climate suitability, and importance of the lineage to the evolutionary legacy of the species as outlined in our study, warrants full or partial consideration. The lineage could be important for understanding the resilience of this species as well as a possible trajectory of trailing edge populations in the northerly ranges. It could help address what might occur when climate shifts from an area but a population remains (87). The distinctiveness of the leading edge lineage is more likely a phenomenon of drift from occasional founding events from the core range (90). The importance of the leading edge lineage is for continued and future exchange of locally adapted genes (e.g., cold tolerance and dispersal-promoting) as migrants serially colonize from the core (10). Similarly, the core range could serve as a reservoir for warmer adapted genes with the potential to spread into the leading edge range (72). Our results suggest that prior to the Anthropocene, *E. maculatum* has been well-suited to shifting its range to follow geographic changes in climate, a key quality towards avoiding climate driven extinction (18). But the uncertainty of anthropogenic pressures in more localized, drought-prone, regions could still threaten the persistence of local populations (20). In particular, over-grazing could reduce vegetation that supports their prey base. However, on the north end of the range, in British Columbia, wildfires are of increasing frequency and expected to result in forest conversion to dry grassland ecosystems, especially at lower elevations in areas of rocky mountainous terrain (91). This suggests that suitable habitat for *E. maculatum* in British Columbia and other areas of the Pacific Northwest (92) may increase with climate change in areas where suitable rock/cliff features exist. Much of the finer scale, contemporary, genetic processes of range expansion for the species are still unknown. However, an accumulation of localized extirpations in the core range could potentially slow the rate of beneficial alleles spreading toward the leading edge (77,90,93).

## Acknowledgements

We thank the New Mexico Museum of Natural History and Science, Museum of Southwestern, Burke Museum, University of Washington, Slater Museum of Natural History, University of Puget Sound, and Royal Ontario Museum for historical tissue loans. Thanks to Brad Butterfield for instruction on ecological niche modeling. This material is based upon work supported by the National Science Foundation Graduate Research Fellowship under Grant No. (NSF) 1938054.

## Supporting information

### S1 Supporting Information

Maxent continuous projection maps, performance metrics, and predictor importance.

